# Many dissimilar protein domains switch between α-helix and β-sheet folds

**DOI:** 10.1101/2021.06.10.447921

**Authors:** Lauren L. Porter, Allen K. Kim, Swechha Rimal, Loren L. Looger, Ananya Majumdar, Brett D. Mensh, Mary Starich

**Affiliations:** National Library of Medicine, National Center for Biotechnology Information, National Institutes of Health, Bethesda, MD 20894, USA; National Heart, Lung, and Blood Institute, Biochemistry and Biophysics Center, National Institutes of Health, Bethesda, MD 20892, USA; Howard Hughes Medical Institute, Janelia Research Campus, Ashburn, VA 20147, USA; The Johns Hopkins University Biomolecular NMR Center, The Johns Hopkins University, Baltimore, MD 21218, USA

## Abstract

Hundreds of millions of structured proteins sustain life through chemical interactions and catalytic reactions^1^. Though dynamic, these proteins are assumed to be built upon fixed scaffolds of secondary structure, α-helices and β-sheets. Experimentally determined structures of over >58,000 non-redundant proteins support this assumption, though it has recently been challenged by ∼100 fold-switching proteins^2^. These “metamorphic^3^” proteins, though ostensibly rare, raise the question of how many uncharacterized proteins have shapeshifting–rather than fixed–secondary structures. To address this question, we developed a comparative sequence-based approach that predicts fold-switching proteins from differences in secondary structure propensity. We applied this approach to the universally conserved NusG transcription factor family of ∼15,000 proteins, one of which has a 50-residue regulatory subunit experimentally shown to switch between α-helical and β-sheet folds^4^. Our approach predicted that 25% of the sequences in this family undergo similar α-helix ⇌ β-sheet transitions, a frequency two orders of magnitude larger than previously observed. Our predictions evade state-of-the-art computational methods but were confirmed experimentally by circular dichroism and nuclear magnetic resonance spectroscopy for all 10 assiduously chosen dissimilar variants. These results suggest that fold switching is a pervasive mechanism of transcriptional regulation in all kingdoms of life and imply that numerous uncharacterized proteins may also switch folds.

## Main Text

For over 60 years, biological science has been heavily influenced by the protein folding paradigm, which asserts that a protein assumes one fold specified by its amino acid sequence^5^. Fold-switching proteins challenge this paradigm by remodeling their secondary and tertiary structures and changing their functions in response to cellular stimuli^2^. These proteins regulate diverse biological processes^6^ and are associated with human diseases such as cancer^7^, malaria^8^, and COVID-19^9^. Nevertheless, the ostensible rarity of fold switching leaves open the question of whether it is a widespread molecular mechanism or a rare exception to the well-established rule.

A major barrier to assessing the natural abundance of fold-switching proteins has been a lack of predictive methods to identify more. Whereas computational methods for rapid and accurate prediction of secondary and tertiary structure for single-fold proteins have been established^10-12^, methods to simply classify fold switchers have lagged. This comparative lack of progress arises from the small number of experimentally observed fold switchers (<100), hampering the discovery of generalizable characteristics that distinguish them from single folders. As a result, essentially all naturally occurring fold switchers have been discovered by chance^13^.

Here we use a sequence-based approach^14,15^ to assess the prevalence of fold switching in the NusG protein superfamily, the only family of transcriptional regulators known to be conserved from bacteria to humans^16^. Housekeeping NusGs (hereafter called NusGs) exist in nearly every known bacterial genome and associate with transcribing RNA polymerase (RNAP) at essentially every operon, where they promote transcription elongation. By contrast, specialized NusGs (NusG^SP^s), such as UpxY, LoaP, and RfaH, promote transcription elongation at specific operons only^17^. Atomic-level structures of NusG and RfaH have been determined (**Fig. 1a**). They share a two-domain architecture with an N-terminal NGN domain that binds RNAP, and a C-terminal β-roll domain. By contrast, the C-terminal domain (CTD) of *Escherichia coli* RfaH switches between two disparate folds: an α-helical hairpin that inhibits RNAP binding except at operon polarity suppressor (*ops*) DNA sites and a β-roll that binds the S10 ribosomal subunit, fostering efficient translation^4^. This reversible change in structure and function is triggered by binding to both *ops* DNA and RNAP^18^.

**Fig. 1.**
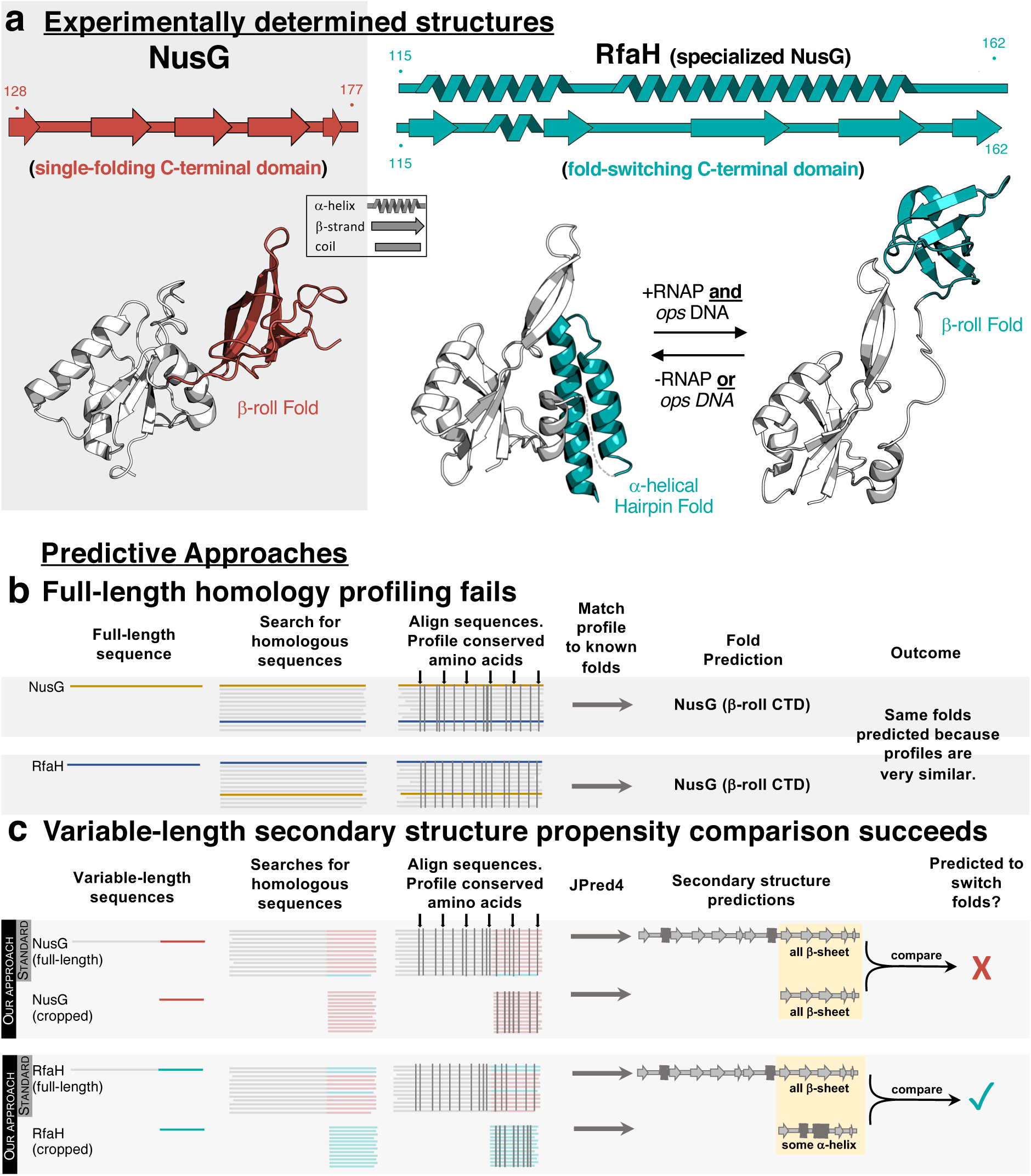
Variable-length secondary structure propensity comparison discriminates between fold-switching RfaH and single-folding NusG. **(a)** Experimentally determined secondary structures and folds of single-folding NusG and the autoinhibited/active NusG^SP^, RfaH (α-helical hairpin/β-roll folds, respectively). Dashed line represents the NTD-CTD linker missing in the RfaH crystal structure. NusG/RfaH CTDs are colored red/teal; NTDs are gray. **(b)**. Profile-based methods fail to identify structural differences between full-length NusG and RfaH because both proteins have similar conservation patterns. Vertical gray bars indicate positions of conserved amino acids. **(c)**. Variable-length secondary structure propensity comparison identifies structural differences between single-folding NusG and fold-switching RfaH. Secondary structure propensities of both the full-length and cropped (CTD) sequences of NusG (above) and RfaH (below) are determined using JPred4. Typically JPred4 is run on full-length sequences only (“standard” in gray box). While both full-length and cropped NusG sequences have similar amino acid conservation patterns (gray vertical lines, top gray panel), conservation patterns differ for full-length and cropped RfaH (gray vertical lines, bottom gray panel). Similar/different full-length and cropped conservation patterns lead to similar/different secondary structure predictions, implying that NusG does not switch folds (top) while RfaH does (bottom).

### Pervasive fold switching is predicted in the NusG superfamily

Profile-based methods identify similar conservation patterns in the amino acid sequences of *E. coli* RfaH and NusG and thus predict that they assume the same folds (**Fig. 1b**). To circumvent this problem, we developed an approach that compares the secondary structure propensities of full-length NusG/NusG^SP^ with those of their CTD sequences. This approach is based on the hypothesis that single-folding and fold-switching NusG CTDs have different secondary structure propensities that can change depending on their molecular context, i.e. whether or not they are tethered to an NTD. Consistent with this hypothesis, JPred4^19^ secondary structure predictions discriminate between the sequences of RfaH and NusG CTDs with experimentally determined atomic-level structures (**Figs. 1c**). These sequence-based calculations consistently indicate that NusG CTD sequences fold into β-strands connected by coils, whereas the *E. coli* RfaH CTD assumes a mixture of α-helix, β-strand, and coil. Thus, our results suggest that JPred4 can distinguish between the sequences of single-folding NusGs and the fold-switching NusG^SP^, RfaH.

To test the generality of our secondary-structure-based approach, we collected 15,516 non-redundant NusG/NusG^SP^ sequences and tested our approach on each hit with a long enough CTD to make predictions (**Methods**), totaling 15,195 sequences. In total, 25% of proteins in the NusG superfamily were predicted to switch folds, a considerable subpopulation with over 3500 sequences.

To estimate the false-negative and false-positive rates of these predictions, we exploited known operon structures of NusG and several specialized homologs^17^ as an orthogonal method to annotate sequences as NusGs or NusG^SP^s. We mapped the sequences used for prediction to sequenced bacterial genomes (**Methods**) and analyzed each sequence’s local genomic environment for signatures of co-regulated genes. Of the 15,195 total sequences, 5,435 mapped to contexts consistent with housekeeping NusG function. Only 30 of these were predicted to switch folds, suggesting a false-positive rate of 0.6% for fold-switch predictions. Performing a similar calculation in 1078 previously identified RfaHs^17^ (**Supplementary Table 1**), 31 were predicted single folders. These results suggest that fold switching is widely conserved among RfaHs, which, if correct, indicates a false positive rate of 3% (31/1078). Of the remaining 8,223 sequences with high-confidence predictions (**Methods**), 30% were predicted to switch folds.

### Experimental validation of fold-switch predictions

A representative group of variants with dissimilar sequences was selected for experimental validation. First, all NusG-superfamily sequences were clustered and plotted on a force-directed graph, hereafter called NusG sequence space (**Figure 2a, Extended Data Fig. 2, Supplementary Table 1**). The map of this space, in which clusters with higher sequence similarity are grouped closer in space, revealed that some putative fold-switching/single-folding nodes cluster together within sequence space (upper/lower groups of interconnected nodes), while other regions had mixed predictions (left/right groups of interconnected nodes). Candidates selected for experimental validation came from distinct nodes, had diverse genomic annotations, and originated from different bacterial phyla (**Extended Data Fig. 3, Supplementary Table 2**).

**Fig. 2.**
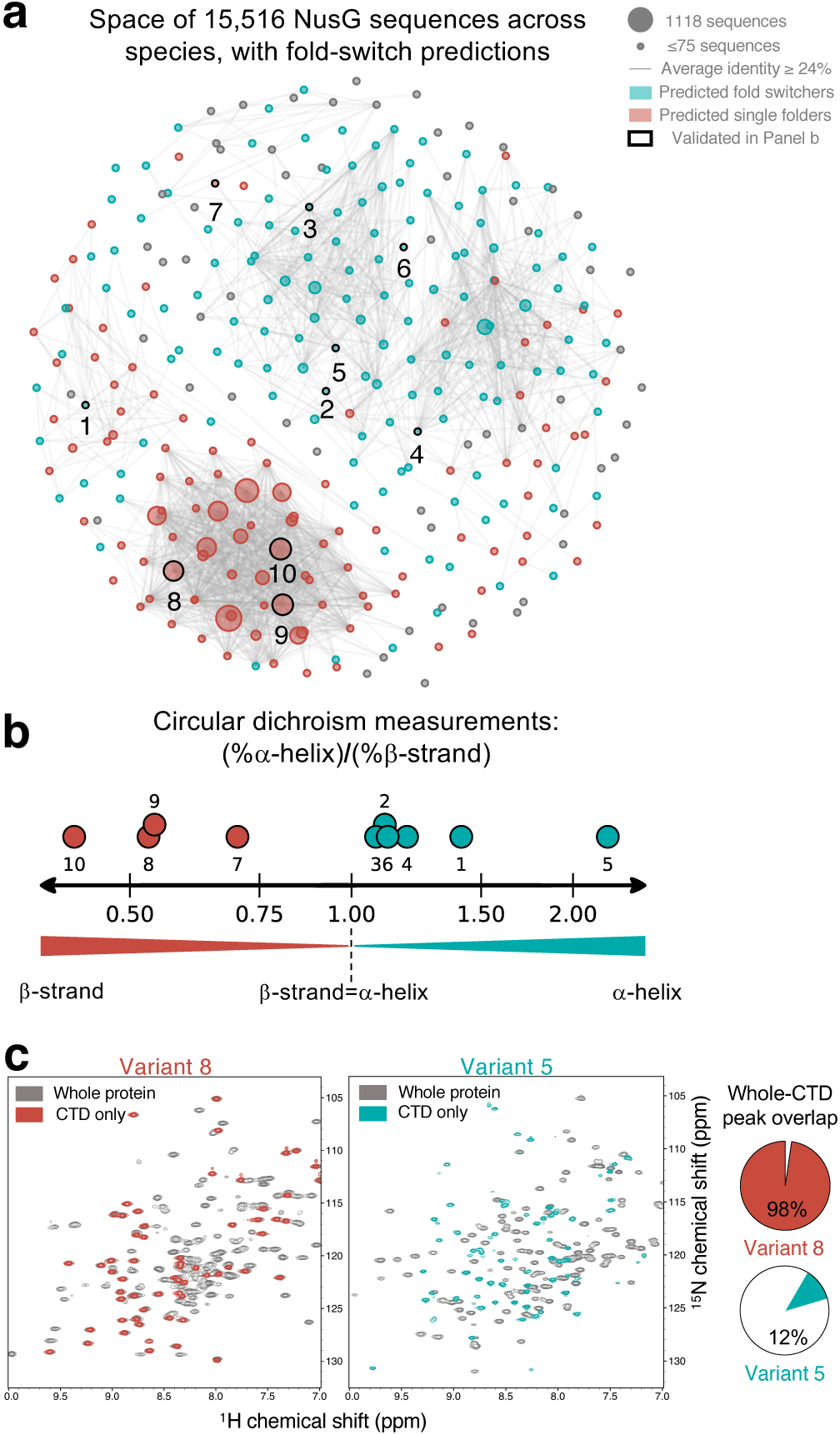
RfaH/NusG sequence space. **(a)**. Force-directed graph of 15,516 full-length RfaH/NusG sequences. The largest node contains 1118 sequences; all nodes with 75 sequences or fewer are the same (smallest) size. Edges connecting the graph represent an average aligned identity between the sequences in two nodes ≥24%. Nodes labeled in teal/red were predicted to be fold switchers/single folders, on average; gray nodes contained only sequences with low-confidence predictions. **(b)**. Fraction of α-helix:β-sheet measured from CD. Dotted line (1.0) represents the minimum α-helix:β-sheet ratio for RfaH-like spectra. All ratios for predicted fold switchers are above the cutoff; all ratios for predicted single folders fall below. Numerical labels shown in **(a)** correspond to variant numbers. Numbers shown on a log_2_ scale. **(c)**. The ^1^H-^15^N HSQCs of full-length and CTD variants of a putative single-folder (Variant #8) are nearly superimposable (98% overlap), while the HSQCs of full-length and CTD variants of a putative fold switcher (Variant #5) differ significantly (12% overlap).

Circular dichroism (CD) spectra of 10 full-length variants were collected. We expected the spectra of fold switchers to have more helical content than single folders because their CTDs have completely different structures (RfaH: all α-helix, NusG: all β-sheet), while the secondary structure compositions of their single-folding NTDs are expected to be essentially identical. *E. coli* RfaH (variant #3) and *E. coli* NusG (variant #9) were initially compared because their atomic-level structures have been determined^20,21^. As expected, their CD spectra were quite different (**Extended Data Fig. 4a**): *E. coli* RfaH had a substantially higher α-helix:β-strand ratio (1.1) than *E. coli* NusG (0.54) – consistent with solved structures (**Fig. 2b, variants #3 and #9**).

All remaining predictions were also consistent with their corresponding CD spectra (**Figure 2b, Supplementary Table 2**). Specifically, five predicted fold switchers had RfaH-like CD spectra, suggesting ground-state helical bundle conformations: two RfaHs (variants #2, #6), a LoaP (variant #1), an annotated NusG (variant #4), and an annotated “NGN domain-containing protein” (variant #5). Furthermore, the remaining three predicted single folders had NusG-like CD spectra: two annotated NusGs (variants #8, #10) and one UpbY/UpxY (variant #7).

We then assessed whether putative fold-switching CTDs could assume β-sheet folds in addition to the α-helical conformations suggested by CD. Previous work^4^ has shown that the full-length RfaH CTD folds into an α-helical hairpin while its isolated CTD folds into a stable β-roll. Thus, we determined the CD spectra of five isolated CTDs: three from putative fold switchers and two from putative single folders. All of them had low helical content and high β-sheet content (**Extended Data Fig. 4b**), strongly suggesting that the CTDs of all three predicted fold switchers can assume both α-helical hairpin and β-roll topologies.

Two variants were then characterized at higher resolution using nuclear magnetic resonance (NMR) spectroscopy. Previous work^4^ has shown that the isolated CTD of RfaH has a significantly different 2D ^1^H-^15^N HSQC spectrum than full-length RfaH, whose CTD folds into an α-helical hairpin. Thus, we conducted similar experiments on one single-folding variant (#8) and one putative fold switcher (variant #5). The backbone amide resonances of the full-length and CTD forms of variant #8 were 98% superimposable, whereas the full-length and CTD forms of variant #5 shared only 7/58 common backbone amide peaks (**Fig. 2c**). This result demonstrates that, as predicted, variant #8 does not switch folds. It is also consistent with the prediction that variant #5 switches folds because large backbone amide shifts indicate either refolding or a large change in protein interface. Both occur in fold-switching RfaH but not in single-folding NusG. Subsequently, assigned backbone amide resonances were used to characterize the secondary structures of both CTD variants at higher resolution (**Extended Data Fig. 4c, Supplementary Table 3**). Both were consistent with the β-roll fold, again suggesting that variant #5 switches folds.

These results, though a very small proportion of the sequences in this superfamily, support the accuracy of our predictions and indicate that:

1. Some, but not all, NusG^SP^s besides RfaH probably switch folds. Specifically, full-length LoaP (variant #1), which regulates the expression of antibiotic gene clusters^22^, had an RfaH-like CD spectrum, whereas full-length UpbY, a capsular polysaccharide transcription antiterminator from *B. fragilis* (variant #7), appears to assume a NusG-like fold.
2. Some annotated NusGs have RfaH-like CD spectra (variant #4), the result of incorrect annotation. Indeed, the genomic environment of variant #4 (**Methods**) suggests that is a UpxY, not a NusG.
3. The fold-switching mechanism appears to be conserved among annotated RfaHs with low sequence identity (≤32%, variants #2, #3, and #6), a possibility proposed previously^23^, though without experimental validation. Also, “NGN domain-containing protein” variant #5 is genomically inconsistent with being a NusG and is likely another RfaH.

To benchmark the performance of our secondary-structure-based approach, we assessed whether machine learning and template-based methods could also distinguish between fold switchers and single folders in the NusG superfamily. Specifically, we tested AlphaFold2^11^, Robetta^24^, EVCouplings^25^, and Phyre2^26^ on variants #1-6, whose CD spectra were all RfaH-like and were thus presumed to switch folds. All methods predicted only one conformation per variant, (**Extended Data Fig. 5**), almost all of which were β-sheet except for AlphaFold2’s predictions of *E. coli* RfaH (variant #3), whose experimentally determined structure^20^ was in its training set, and variant #6, whose sequence is nearest and best connected with *E. coli* RfaH in Sequence Space (**Figure 2a**). All predictions were generated using multiple sequence alignments containing mixtures of RfaH and NusG sequences, and the predicted amino acid contacts from Robetta and EVCouplings corresponded with the NusG-like β-roll fold (**Extended Data Fig. 6**). These results suggest that β-roll couplings present in both single-folding and fold-switching sequences might overwhelm any α-helical couplings unique to fold-switching sequences.

We then performed coevolutionary analysis on a subset of sequences that our approach predicted to switch folds (**Methods**). Specifically, we clustered putative fold switchers by their secondary structure predictions and did coevolutionary analysis on the cluster containing the *E. coli* RfaH sequence using GREMLIN^27^. The residue-residue contacts generated from these sequences differed substantially from the NusG-like couplings generated before (**Figure 3**). Furthermore, GREMLIN couplings calculated from the alignments used by EVCouplings and Robetta corresponded with the β-roll fold only (**Extended Data Fig. 6**), demonstrating that the JPred-filtered sequence alignment—not the GREMLIN algorithm—was responsible for the discovery of alternative contacts.

**Fig. 3.**
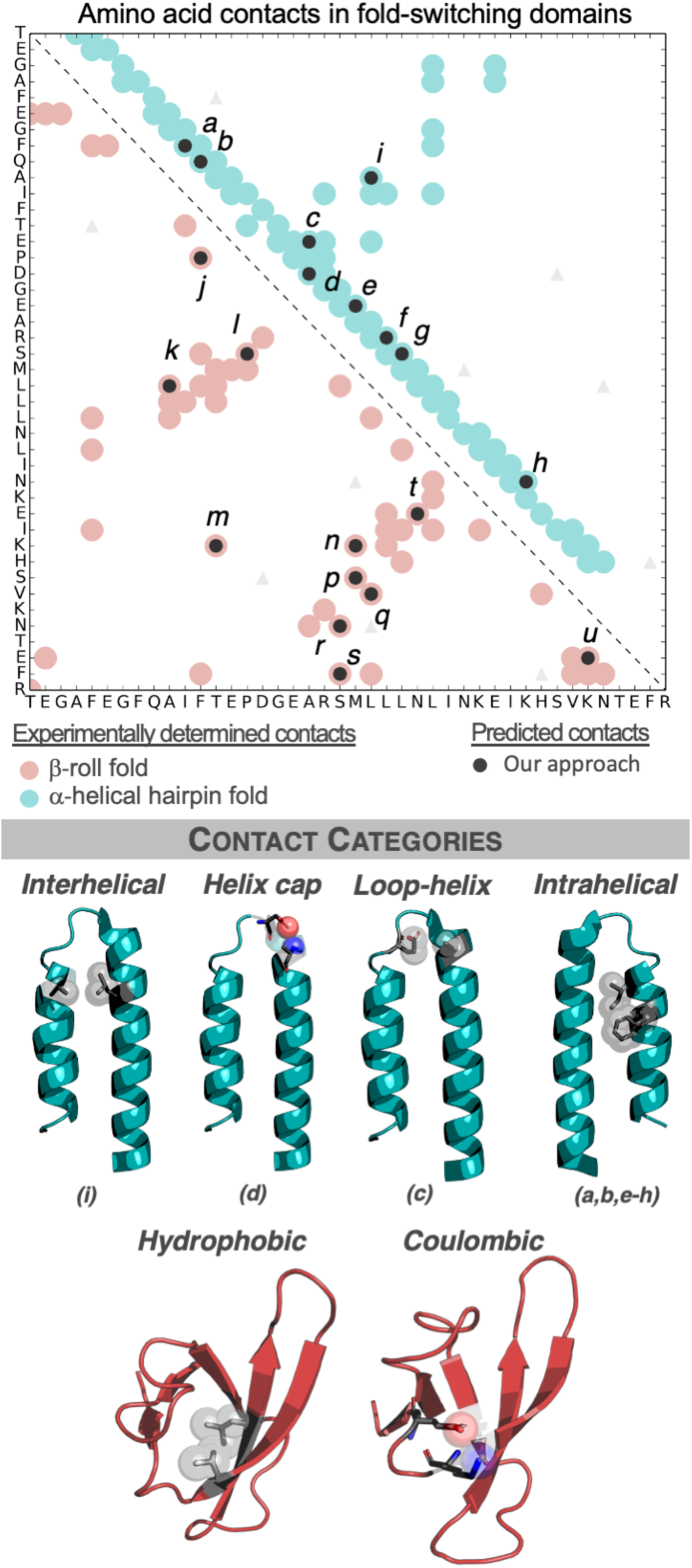
Fold-switching sequences have conserved amino acid contacts from both folds. Predicted amino acid contacts from fold-switching sequences (dark gray circles) correspond to both the β-roll fold (PDB ID: 2LCL, red circles) and the α-helical hairpin fold (PDB ID: 2OUG, chain A, teal circles). Couplings that do not correspond to experimentally observed contacts are shown as light triangles. Categories of amino acid contacts from both folds use the alphabetically labeled contacts in the plot above them.

This analysis of putative fold-switching CTDs indicates evolutionary coupling of residue-residue contacts unique to two distinct folds. For the α-helical fold, six intrahelical hydrophobic contacts and one set each of interhelical contacts, strand-helix contacts, and helix-capping contacts were observed (**Fig. 3**). Overall, 96% of interhelical contacts were hydrophobic, 94% of helix-capping residues could potentially form an i-4→i or i→i backbone-to-sidechain hydrogen bond, 85% of residues in the helix-loop interaction had a charged residue in one position, but not both, and 80% of residues in intrahelical contact *a* were both hydrophobic. The remaining contacts gave more mixed results, perhaps due to hydrophobic residues contacting the hydrophobic portion of their hydrophilic partners. Contacts from the β-roll fold, identified by both GREMLIN and EVCouplings/Robetta, were categorized as Coulombic and hydrophobic **(Extended Data Fig. 7)**. Previous work has shown that interdomain interactions also contribute significantly to RfaH fold switching^4^. Unfortunately, these interactions could not be identified by coevolutionary analysis (**Extended Data Fig. 8**), a likely result of the limited number of JPred-filtered sequences available.

### Fold-switching CTDs are diverse in sequence, function, and taxonomy

It might be reasonable to expect fold-switching CTD sequences to be relatively homogeneous, especially since variants of another fold switcher, human XCL1, lose their ability to switch folds below a relatively high identity threshold (60%)^28^. The opposite is true. Sequences of putative fold-switching CTDs are significantly more heterogeneous (20.4% mean/19.4% median sequence identity) than sequences of predicted single folders (40.5% mean/42.5% median sequence identity, **Fig. 4a**). Accordingly, among the sequences tested experimentally, similar mean/median sequence identities were observed: 21.0%/21.1% (fold switchers), 43.2%/41.2% (single folders, **Fig. 4b**). Furthermore, fold-switching CTDs were predicted in most bacterial phyla, and many were predicted in archaea and eukaryotes as well (**Fig. 4c, Supplementary Table 4**). These results suggest that many highly diverse CTD sequences can switch folds between an α-helical hairpin and a β-roll in organisms from all kingdoms of life.

**Fig. 4.**
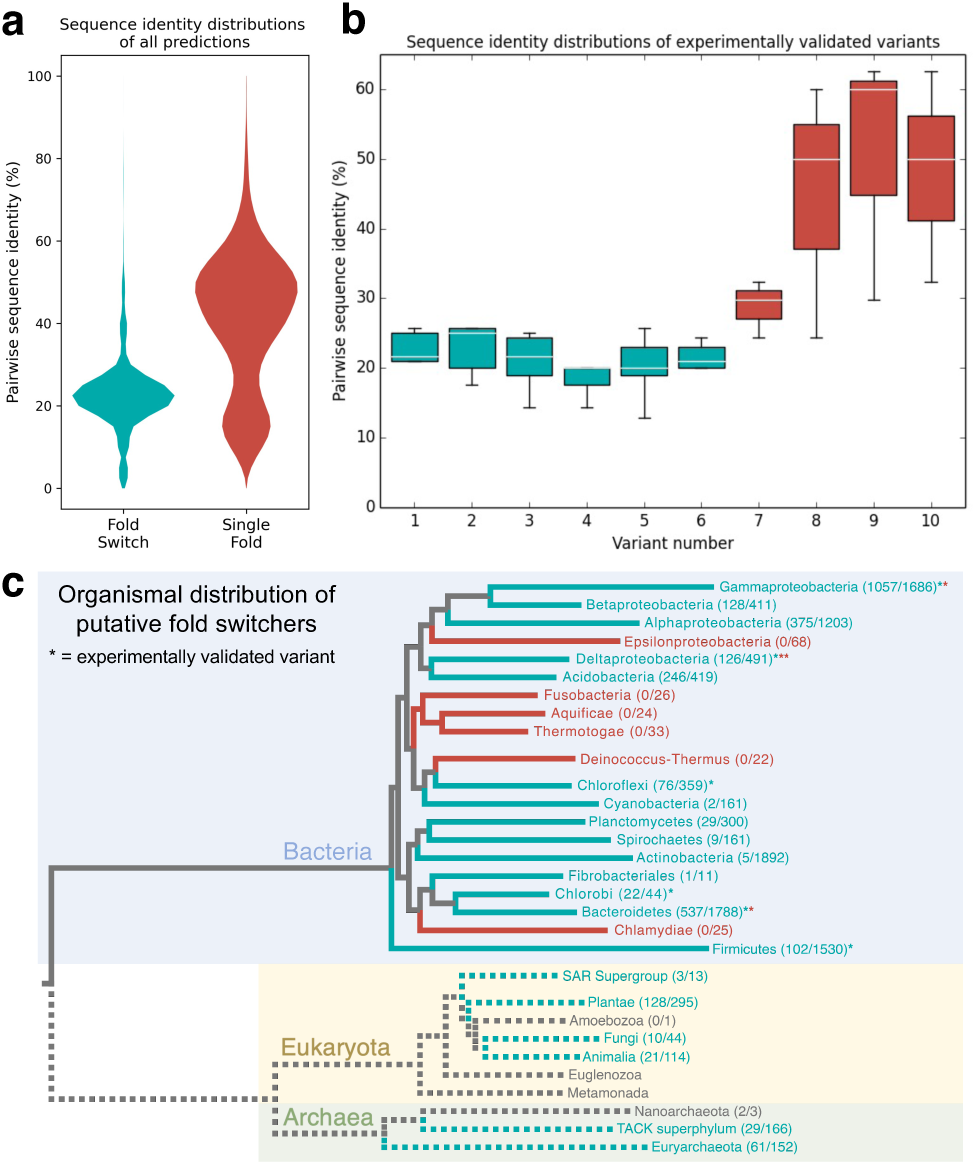
The sequences of fold-switching CTDs are highly diverse and found in a wide variety of bacterial phyla. **(a)** Violin plots of pairwise sequence identities differ significantly for putative fold switchers and putative single folders. On average, pairwise sequence identities are lower for putative fold switchers (20.4%) than single folders (40.5%). **(b)** Box-and-whiskers plots of pairwise sequence identities of fold-switching and single-folding CTDs of variants 1-10 in **Figure 2**. These distributions are consistent with the violin plots in panel **a. (c)** Fold-switching CTDs are predicted in many bacterial phyla and other kingdoms of life. Numbers next to taxa represent #predicted fold switchers/#total sequences. Gray branches represent unidentified common ancestors, since the evolution of fold-switching NusGs is unknown. Dotted lines represent lower-confidence predictions since fold switching has not been confirmed experimentally in archaea and eukaryota. Fold-switching/single-folding predictions are represented by teal/red colorings; predictions in branch3e5s with fewer than 10 sequences are gray.

Why might the sequence diversity of fold-switching CTDs exceed those of single folders? Functional diversity is one likely explanation^29^. Previous work has shown that NusG^SP^s drive the expression of diverse molecules from antibiotics to toxins^17^. Our approach suggests that many of these switch folds. Furthermore, since helical contacts are conserved among at least some fold-switching CTDs, it may be possible that CTD sequence variation is less constrained in other function-specific positions. The fold-switching mechanism of RfaH allows it to both regulate transcription and expedite translation, presumably quickening the activation of downstream genes. Fold-switching NusG^SP^s are likely under strong selective pressure to conserve this mechanism when the regulated products control life-or-death events, such as the appearance of rival microbes or desiccation. Supporting this possibility, NusG^SP^s usually drive operons controlling rapid response to changing environmental conditions such as macrolide antibiotic production^22^, antibiotic-resistance gene expression^17^, virulence activation^30^, and biofilm formation^31^.

Our approach was sensitive enough to predict fold-switching proteins, setting it apart from other state-of-the-art methods. These other methods assume that all homologous sequences adopt the same fold, as evidenced by their use of sequence alignments that contained both fold-switching and single-folding sequences. These mixed sequence alignments biased their predictions. While those predictions are partially true since both fold-switching and single-folding CTDs can fold into β-rolls, they miss the alternative helical hairpin conformation and its regulatory function^16^. Computational approaches that account for conformational variability and dynamics, a weakness in even the best predictors of protein structure^11^, could lead to improved predictions. This need is especially acute in light of recent work showing how protein structure is influenced by the cellular environment^32^, and it could inform better design of fold switchers, a field that has seen limited success^33-35^.

Our results indicate that fold switching is a pervasive, evolutionarily conserved mechanism. Specifically, we predicted that 25% of the sequences within a ubiquitous protein family switch folds and observed coevolution of residue-residue contacts unique to both folds. This sequence-diverse dual-fold conservation challenges the protein folding paradigm and indicates that foundational principles of protein structure prediction may need to be revisited.

The success of our method in the NusG superfamily suggests that it may have enough predictive power to identify fold switching in protein families where only single folders have been observed. Such predictions would be particularly useful since many fold switchers are associated with human disease^6-9^. Given the unexpected abundance of fold switching in the NusG superfamily, there may be many more unrelated fold switchers to discover.

## Acknowledgements

L.L.P. thanks Marius Clore and Carolyn Ott for constructive discussions. We also thank George Rose, Liskin Swint-Kruse, Gisela Storz, Juan Bonifacino, Nico Tjandra, David Nyenhuis, and Daniel Morris for helpful comments concerning the text and Drs. Juliette Lecomte and Christos Kougentakis for helping us to collect NMR data at the Johns Hopkins University Biomolecular NMR Center. This work utilized resources from the NHLBI Biophysics Core, the NHLBI Protein Expression Facility, and the NIH HPS Biowulf cluster (http://hpc.nih.gov), and it was supported in part by the Intramural Research Program of the National Library of Medicine, National Institutes of Health and Howard Hughes Medical Institute.

## Author contributions

Conceptualization: LLP, LLL

Methodology: LLP, LLL, AK, MS, AM

Software: LLP, LLL, AK

Investigation: LLP, LLL, AK

Data Curation: LLL, LLP

Visualization: LLP, BDM, AK

Writing – original draft: LLP

Writing – review & editing: LLP, BDM, LLL, MS, AK

Supervision: LLP, LLL

Project administration: LLP

Funding acquisition: LLP, LLL

## Data and materials availability

All data and code can be found at: https://github.com/porterll/sequence_space. Constructs for protein expression are available upon request.

## Methods

### Identification of NusG-like sequences

NusG-like sequences were identified from the October 2019 Uniprot90^36^ database using an iterative BLAST^37^ approach. Specifically, the *E. coli* RfaH sequence (Uniprot ID Q0TAL4) was BLASTed against the database. All hits with a maximum e-value of 10^−4^ were aligned using Clustal Omega^38^, which generated their sequence identity matrices from the resulting alignment. Sequences were clustered by their identities using the agglomerative clustering algorithm from the python module scikit-learn^39^. Sequence identity between proteins in each cluster was ≥ 78%. Randomly selected sequences from the 25 largest clusters were then individually BLASTed against the Uniprot90 database, and the resulting hits were combined; redundant identical hits from independent searches were removed. This procedure (search-align-cluster) was repeated two additional times to generate the full list of 15,516 sequences in 305 clusters.

### Determination of CTDs

Sequences of annotated RfaHs were aligned to the sequence of *E. coli* RfaH (Uniprot ID Q0TAL4) using Clustal Omega^38^. CTDs were defined as up to 50 residues, but not shorter than 40 if the CTD region comprised <50 residues, beginning with the positions that aligned to the RfaH sequence *KVIIT*. Sequences of proteins not annotated as RfaH were aligned to the *E. coli* NusG sequence (Uniprot ID P0AFG0) using Clustal Omega. CTDs were defined as 50 residues beginning with positions that aligned the NusG sequence *EMVRV*. Because of their diversity, sequences from each individual cluster were aligned against the NusG sequence separately, each using Clustal Omega. The number of sequences with CTDs long enough to make these predictions totaled 15,195 (**Supplementary Table 1**), 98% of all NusG-like sequences identified.

### JPred4 predictions

JPred4^19^ predictions were carried out as in^14^, sections 2.4 and 2.6. In further detail, they were first performed on all 50-residue CTD sequences using two databases: the JPred database (http://www.compbio.dundee.ac.uk/jpred/about_RETR_JNetv231_details.shtml) from 2014 and the Uniprot90 database from January 2021. Sequences of each prediction were aligned against the *E. coli* NusG sequence (beginning with *EMVRV*) using Biopython^40^ Bio.pairwise2.localxs with gap opening/extension scores of -1.0/-0.5. Secondary structure predictions of the sequence in question and of *E. coli* NusG were reregistered according to the resulting pairwise alignments and compared as in^14^. Predictions were considered high-confidence if at least 5 sequences were in the MView^41^-generated alignments used by JPred.

We found that the first 10 residues in these 50-residue sequences were similar enough to NusG CTDs that NusG-like sequences overwhelmed sequence alignments informing the predictions, and many likely fold-switching sequences were predicted to be single folders. To circumvent this problem, predictions from both databases were rerun on 40-residue sequences (starting with the first residue that aligned to *ADFNG…* for NusG sequences and *FQAIF*… for RfaH sequences). Predictions were made as with 50-residue sequences. All predictions reported in the main text were from 40-residue sequences, except those in **Fig. 1b**.

### Force-directed graph

The 305 clusters generated from all full-length NusG sequences were plotted on a force-directed graph using the *spring_layout* function from python NetworkX^42^ with a spring constant of 0.3 and 1000 iterations. Nodes with ≥50% of sequences predicted to switch folds were colored teal; nodes with <50% of sequences predicted to switch folds were colored red. Nodes with no predictions were colored gray. Nodes 1 and 7 were colored differently from their average predictions (single folding, Node 1; fold-switching, Node 7) to highlight the prediction of the sequence validated experimentally, which differed from the average. Edges represented average pairwise identities between nodes ≥24%, a threshold taken from^43^ for sequences of 162 residues (the length of *E. coli* RfaH).

### Genomic analysis of sequences

The annotated genomes (protein .fasta and .gtf annotation) of 31,554 bacterial species were downloaded from Ensembl Bacteria in April 2021. Genomic annotation of NusG was defined as being within 10 kb of a gene annotated as either “SecE,” “RplK,” “RplA,” or “ribosomal protein L11” by text matching. Most bacterial genomes are incompletely assembled and annotated – the genes were required to be within the same chromosome, contig, or plasmid. Each Uniprot sequence in the database of 15,516 was mapped to an Ensembl locus if the species was consistent, and if sequence identity was greater than 90%. Annotation was fetched from Ensembl, as well – this was usually, but not always, consistent with the Uniprot annotation.

Of the 15,516 Uniprot sequences, 7975 mapped to Ensembl genomes. Cursory analysis of some non-mapping sequences suggested that: 1) some Ensembl genomes had incomplete collation of all ORFs, and 2) there were frame-shifts and other errors in some Uniprot sequences and some Ensembl genomes. This was also the case for some of the sequences predicted to potentially be fold-switching NusGs: for instance, Uniprot entry A0A0T8ANM4 is frame-shifted relative to the Ensembl genome, producing a C-terminal sequence predicted to switch folds.

Of the 5,435 sequences that mapped to Ensembl loci with *SecE*/*RplK*/*RplA* within 10kb, only 22 had a separation of >1kb, and only 59 had a separation of >270bp – this set of 59 includes 4 proteins predicted to be fold-switching, one of which is a verified RfaH from ^17^, indicating that a shorter threshold of distance to *SecE*/*RplK*/*RplA*, perhaps coupled with determining distance several other conserved *NusG*-*SecE* operon genes, could reduce the false-positive rate caused by mistakenly annotating NusG^SP^s as housekeeping NusGs.

For a small number of sequences that mapped to qualitatively dissimilar genes (e.g., one genomically consistent as being a NusG, another not), the 2^nd^ mapping is given in **Data S1**, beginning in column AH.

Additionally, of the 600 RfaH sequences that mapped to an annotated Ensembl locus, only one fell within a NusG-like operon (∼7kb away).

### Expression and purification of variants 1-16

All variants were ordered from IDT as gBlocks. Except for variant #8, these variants were digested with HindIII and EcoRI and ligated into the pPAL7 vector (Bio-Rad) with an N-terminal 6-His tag cloned using a Q5 mutagenesis kit (New England Biolabs). Variants were transformed into *E. coli* BL21-DE3 cells (New England Biolabs), grown in LB at 37° to an OD_600_ of 0.6-0.8, after which they were incubated at 20°C for 30 minutes, induced with 0.1 mM IPTG, and grown overnight, shaking at 225-250 rpm. Variant #8 was cloned into the same vector as the other variants using In-Fusion and expressed as the other variants but at 18°C instead of 20°C. The cells from all cultures were pelleted at 10,000*xg* for 10 minutes at 4°C, resuspended in 2 mL lysis buffer (50 mM Tris, 150 mM NaCl, 5% glycerol, 1 mM DTT, 10 mM imidazole, pH 8.7) and frozen at -80°C for later purification. Sequencing of all variants was verified by Psomagen.

Thawed cell pellets were resuspended in 25 mL lysis buffer per 1 L of culture grown. 100 mg of DNAseI, 5 mM CaCl_2_, 5 mM MgSO_4_ and 1/2 of a cOmplete EDTA-free protease cocktail inhibitor tablet (Roche) were added per 25 mL of lysis buffer. Cells were lysed by 2 passes through an EmulsiFlex-C3 homogenizer (Avestin). The homogenized lysate was centrifuged for 45 minutes at 40,000*xg* at 4°C, and its soluble fraction was loaded immediately onto either a 1 mL Ni column (GE HisTrap HP) or an Econo-Pac (Bio-Rad) gravity column with 0.5-1 mL IMAC Ni Resin (Bio-Rad). Soluble lysate was loaded on ice for the HisTrap column, and gravity columns were loaded and kept at 4°C. The HPLC Ni columns were washed with 100 mM phosphate and 500 mM NaCl, pH 7.4, equilibrated in 100 mM phosphate, pH 7.4, and eluted by gradient with 0.5 M imidazole, 100 mM phosphate, pH 8.0 at 2 mL/minute on an ÄKTA Avant. The gravity columns were washed and equilibrated with 10 column volumes each of the same buffers, and protein was eluted at 3 different imidazole concentrations: 100 mM, 500 mM and 2M, all in 100 mM phosphate, pH 7.4.

Nickel-purified samples were then loaded onto 1-or 5-mL Profinity eXact^44^ columns (Bio-Rad), washed twice with one column-volume of 2M NaOAc, and eluted with 100 mM phosphate, 10 mM azide, pH 7.4 at 0.2 mL/minute. Cleavage kinetics for some variants (1, 4, and 6) were too slow to get adequate tagless protein. In these cases, columns were equilibrated with 100 mM phosphate, 10 mM azide, pH 7.4 overnight at 4°C. Tagless protein was concentrated in 10 kDa MWCO concentrators (Millipore), and the buffer was exchanged to 100 mM phosphate, pH 7.4. A small amount of high-molecular-weight impurity (<10% of the sample) from variants #1 and #4 was removed by running the tagless sample through a 50 kDa MWCO concentrator (Millipore) and keeping the low molecular weight fraction that passed through the filter. Sample purities were assessed by gel electrophoresis (Thermofisher NuPAGE 4-12% Bis-Tris gels, Thermofisher MES buffer, Bulldog Bio Coomassie Stain), and concentrations were measured on a NanoDrop OneC (Thermo Scientific).

### Circular dichroism (CD) spectroscopy

All CD spectra were collected on Chirascan spectrometers (Applied Photophysics) in 1 mm quartz cuvettes in 100 mM phosphate, pH 7.4. Protein concentrations ranged from 8-12 mM, and scan numbers ranged from 5-10. Scans were averaged, and averaged baselines of buffer-blank 1 mm cuvettes were subtracted from the spectra. The resulting spectra were converted to units of Molar Residue Ellipticity [*θ*]_MRE_ using the following formula:

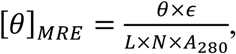

where *θ* is the ellipticity measured by the instrument, *ϵ* is the extinction coefficient determined by Expasy Protparam^45^ (https://web.expasy.org/protparam/), *L* is the path length of the cuvette, N is the number of amino acids, and *A*_280_ is the absorbance at 280 measured by a Nanodrop One (Thermo Scientific). Resulting spectra were entered into the BestSel^46^ webserver (https://bestsel.elte.hu/index.php), so that their ratio of helix (helix+distorted helix):strand (parallel+antiparallel) could be computed.

### CTDs of variants #5 and #8

Full-length variants #5 and #8 were shortened to 64 and 68 residues, respectively, using Q5 mutagenesis (New England Biolabs). Their sequences were confirmed by Sanger sequencing (Psomagen) and are reported in **Supplementary Table 2**.

### Expression and purification of NMR samples

Based on the protocols in^47-49^, BL-21 DE3 cells (New England Biolabs) expressing all NMR samples were grown in LB to an OD_600_ of 0.6 and pelleted at 5000*xg* for 30 minutes at 4°C. The pellets were resuspended in 1X M9 at half of the initial culture volume and pelleted at 5000*xg* for 30 minutes at 4°C. Pellets were then resuspended at ¼ initial culture volume in 2X M9, pH 7.0-7.1, 1 mM MgSO_4_, 0.1 mM CaCl_2_, with 1 g ^15^NH_4_Cl/L, and 4 g of either unlabeled or ^13^C-labeled glucose (Cambridge Isotope Laboratory)/L and equilibrated at 20°C for 30 min, shaking at 225 rpm, then induced with 1 mM IPTG and grown overnight. Cells were pelleted at 10,000xg for 10 minutes at 4°C. All labeled variants were purified by FPLC (ÄKTA Avant 25) using the same methods as variants 1-16 above in 5 mL HisTrap HP columns (Cytiva) and 5 mL Profinity eXact columns (BioRad).

### ^1^H-^15^N HSQCs of variants #5 and #8

All spectra were collected on Bruker Avance II 600 MHz spectrometers equipped with z-gradient cryoprobes and processed with NMRPipe^50^. Variant #8 (full-length and CTD) and variant #5 CTD HSQCs were collected in 100 mM phosphate, pH 7.4 at 298 K. Under those conditions the spectrum of full-length variant 5 was broad, even with 1 mM DTT added, but peaks narrowed upon changing the buffer conditions to 25 mM HEPES, 50 mM NaCl, 5% glycerol, 1 mM DTT, pH 7.5, and collecting the spectrum at 303K. Protein concentrations ranged from 100-300 μM.

### Assignments of KCTD and TCTD

^13^C-labeled 5CTD and 8CTD were expressed and purified as above. For each variant, HNCACB, CBCA(CO)NH, and HNCO experiments were collected on Bruker Avance II 600 MHz spectrometers with cryoprobes. Spectra of 8CTD (80 μM) were collected using nonuniform sampling and were processed with SMILE^51^. All NMR spectra were processed using NMRpipe^50^. Resonances were assigned manually with NMRfam Sparky^52^, and secondary structures were determined using TALOS+^53^.

### Coevolutionary analysis

Structure predictions of the 6 fold-switching variants were calculated by entering their full-length sequences (**Supplementary Table 5**) into the EVCouplings^25^, Robetta^24^, and Phyre2^26^ webservers (https://evcouplings.org,https://robetta.bakerlab.org, http://www.sbg.bio.ic.ac.uk/phyre2/html/page.cgi?id=index). EVCouplings predictions with the recommended e-value cutoffs for chosen: (Variant 1: e-3, 2: e-5, 3: e-5, 4: e-20, 5: E-5, 5: e-5). High-confidence predictions for shorter sequences of 40 or 50 residues could not be obtained from either EVCouplings or Robetta. Predicted residue-residue contacts of *E. coli* RfaH from EVCouplings/Robetta with probabilities ≥ 99%/92% were plotted in **Extended Data Fig. 4a&b**, and residue-residue contacts from GREMLIN^27^ with probabilities ≥90% were plotted in **Fig. 3**.

These thresholds were determined by maximizing the ratio of true positives to false positives. True positives were considered to be couplings with heavy atoms within 5.0 Å in either the 2OUG or the 2LCL crystal structures where at least one of the 2 heavy atoms was from a side chain; one additional contact between residues 140 and 151 was added because they were separated by 5.2 Å within the NMR structure and therefore likely within error of 5.0Å. Contacts were considered hydrophobic if both atoms in contact were hydrophobic, Coulombic if two atoms in contact had opposite charge and C-N-O/C-O-N angles ≥ 90°, and helix caps if the distance between sidechain donor/acceptor ≤4° and C-N-O/C-O-H angles ≥ 90°^54^. All distances and angles were calculated using LINUS^55^.

CTD sequences for GREMLIN webserver (http://gremlin.bakerlab.org/submit.php) analysis in **Fig. 3** were obtained by clustering all JPred predictions by Affinity Propagation using the python Scikit-learn module^39^ with damping of 0.99 and a maximum number of 10,000 iterations. Affinities were precomputed by comparing each 40-residue prediction position-by-position, with the following scores: identical predictions (EE,HH,--): 0, coil:secondary structure discrepancies (H-,E-,-H,-E): 0.5, and helix:strand discrepancies (HE,EH): 10, and selecting the cluster with the sequence of *E. coli* RfaH (639 sequences). These sequences were aligned with Clustal Omega and inputted into GREMLIN. 4 iterations of HHBlits^56^ were run on the initial alignment with E-values of 10^−10^. Coverage and remove gaps filters were both set to 75.

GREMLIN webserver analyses were run on EVCouplings and Robetta multiple sequence alignments seeded with the sequence of *E. coli* RfaH. These alignments were taken from EVCouplings *align* and Robetta .msa.npz files. No additional iterations of HHPred were run on either alignment. Coverage and remove gaps filters were both set to 75.

### Pairwise sequence identities

Pairwise sequence identity matrices of predicted fold-switching/single-folding CTDs were calculated using Geneious. The alignments for these sequences were first manually curated to remove sequences that did not align well with the majority; manually curated alignments retained at least 98% of all sequences. The mean/median sequence identities of these two groups were determined from the upper triangular matrices of each matrix, excluding positions of identity, using numpy^57^. Pairwise sequence identity matrices of the CTDs of the 10 variants were determined with Clustal Omega.

### Phylogenetic tree

The tree in **Figure 4C** was generated by downloading the Interactive Tree of Life^58^ (https://itol.embl.de/itol.cgi), loading it into FigTree^59^, and collapsing branches at the phyletic level, except for Proteobacteria, which were left at the class level because of recent phylogenetic work on proteobacterial RfaH^17^.

Bacterial species from each NusG sequence were obtained from their Uniprot headers. These species were mapped to their respective phyla using TaxonKit^60^ and matched with their predictions. Phyla with fold-switching/single-folding predictions were listed using a python script, and branches of the tree were then colored manually in Adobe Illustrator. Experimentally validated variants from two phyla did not show on the dendrogram in Figure 4: Candidatus Kryptonia and Deferribacteres. They were grouped with Bacteroidetes and Deltaproteobacteria, respectively, their nearest neighbors^61,62^ shown in the tree.

Eukaryotic and archaeal NusG homologs were obtained by running 3 rounds of PSI-BLAST on the nr database with the following seed sequences: L1IE32, A0A0N95N5M7, UPI0005F5777A, A0A2E6HKN0. Redundant sequences were removed using CD-HIT^63^ at a 98% sequence identity threshold (at least 1 amino acid difference).

## Figures

Figures 1C, 2A, 2B, 3A, 3B, 4A, and 4B were generated using Matplotlib^64^. The figures of all protein structures (Figures 1A and 3C) were generated using PyMOL^65^.

**Extended Data Fig. 1.**
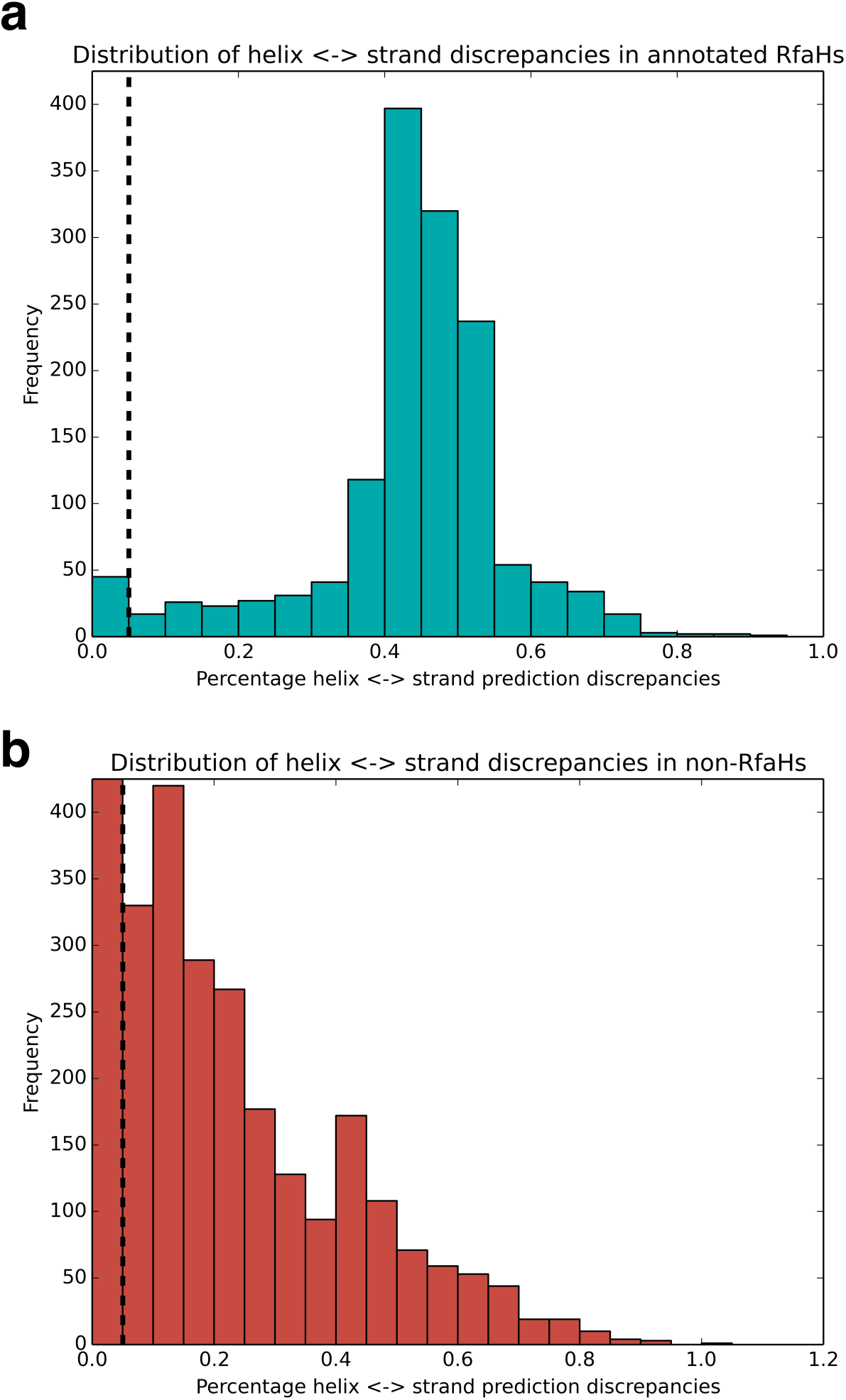
Distributions of prediction discrepancies for Uniprot-annotated RfaHs (a) and sequences not annotated as RfaH (b). Black dotted line is the cutoff point for fold-switching and single-folding predictions: predictions with ≥5% discrepancy to the *E. coli* NusG sequence were predicted to switch folds. This cutoff was taken from Kim, et al. ^14^. Because of the low threshold, experiments were performed on constructs just above/below the threshold (Constructs 1, 4, and 7, respectively, **Supplementary Table 1, Fig. 2**). For comparison, the y-axis of both plots was limited to 410 counts. All bins fell below that threshold except for the first bin of panel B, which had 11,032 counts (and thus is not shown to scale).

**Extended Data Fig. 2.**
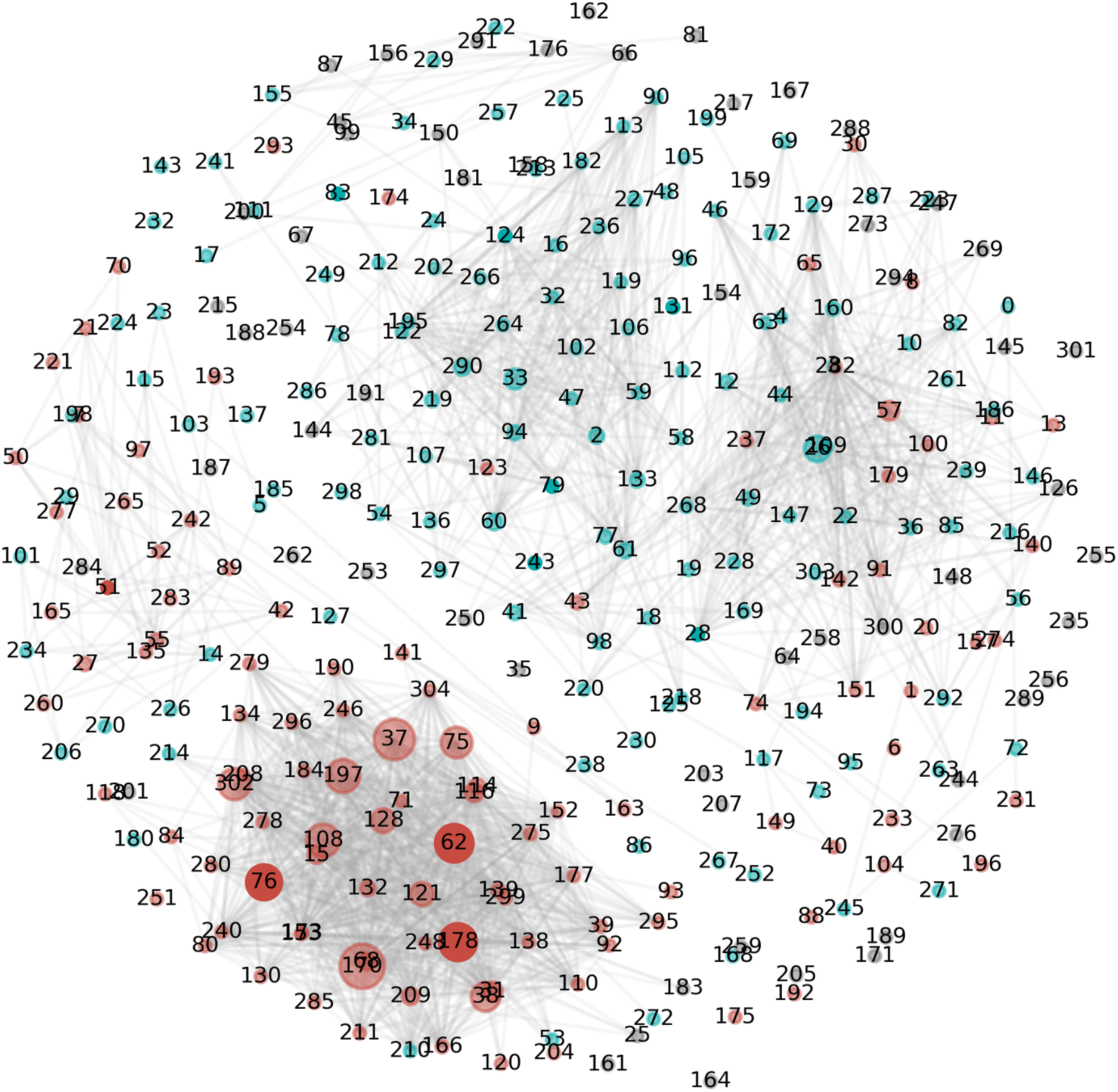
Sequence space diagram with cluster numbers labeled. Numbers correspond to the Cluster IDs (column 3) in **Supplementary Table 1**.

**Extended Data Fig. 3.**
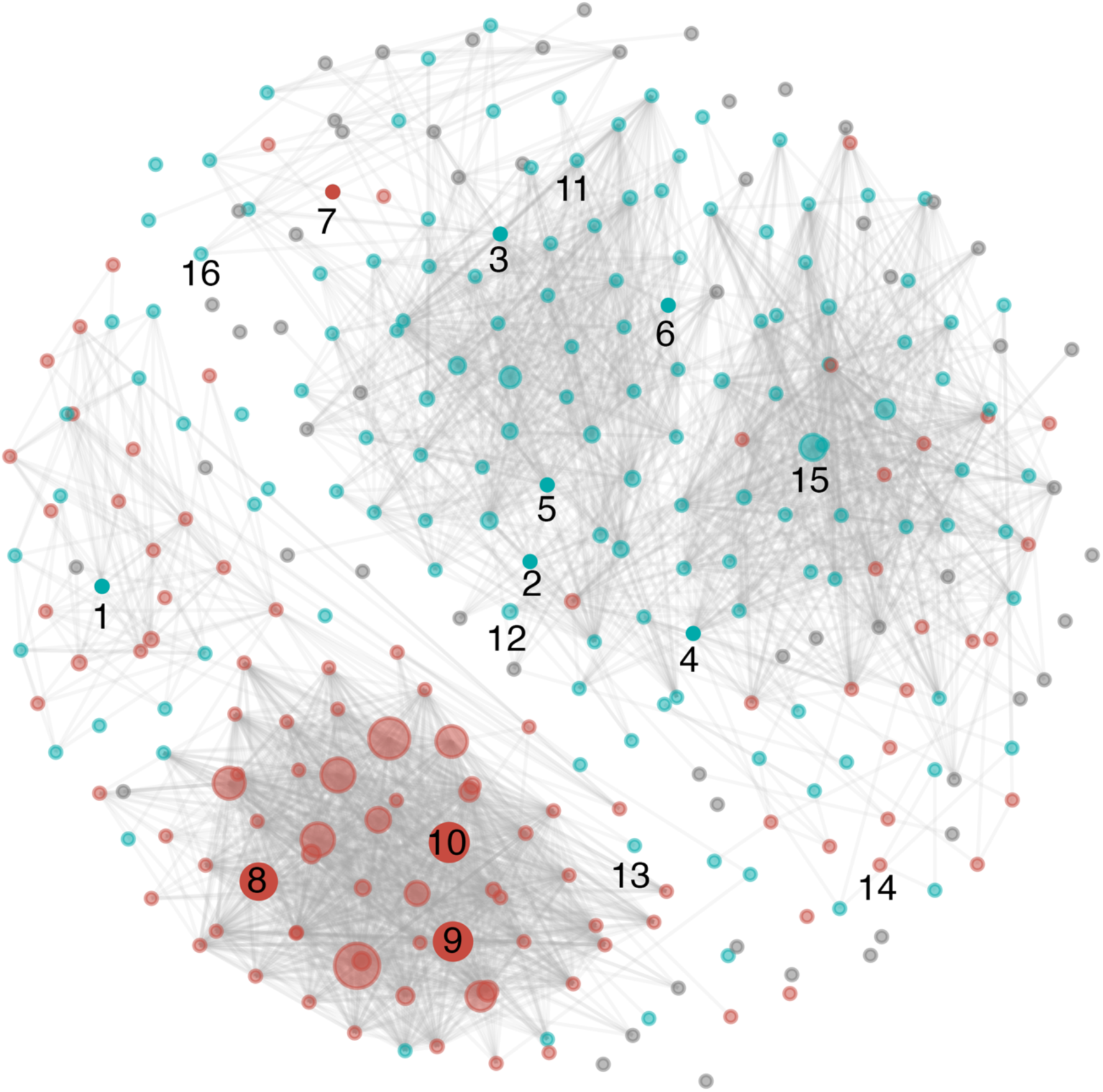
All sequence space constructs tested. Constructs 1-10 are labeled as in Figure 2. Constructs 11-16 were tested, but CD spectra could not be obtained because they did not express (Constructs 11-13 and 15-16) or because they were insoluble (Construct 14). With the exceptions of Constructs 8-10, all labels are directly below the nodes from which sequences were selected. Teal/red nodes: predicted to/not to switch folds on average; no high-confidence predictions were made for gray nodes. More information about each construct can be found in **Supplementary Table 2**. Nodes 1 and 7 were colored differently from their average predictions (single folding, Node 1; fold-switching, Node 7) to highlight the prediction of the sequence validated experimentally, which differed from the average.

**Extended Data Fig. 4.**
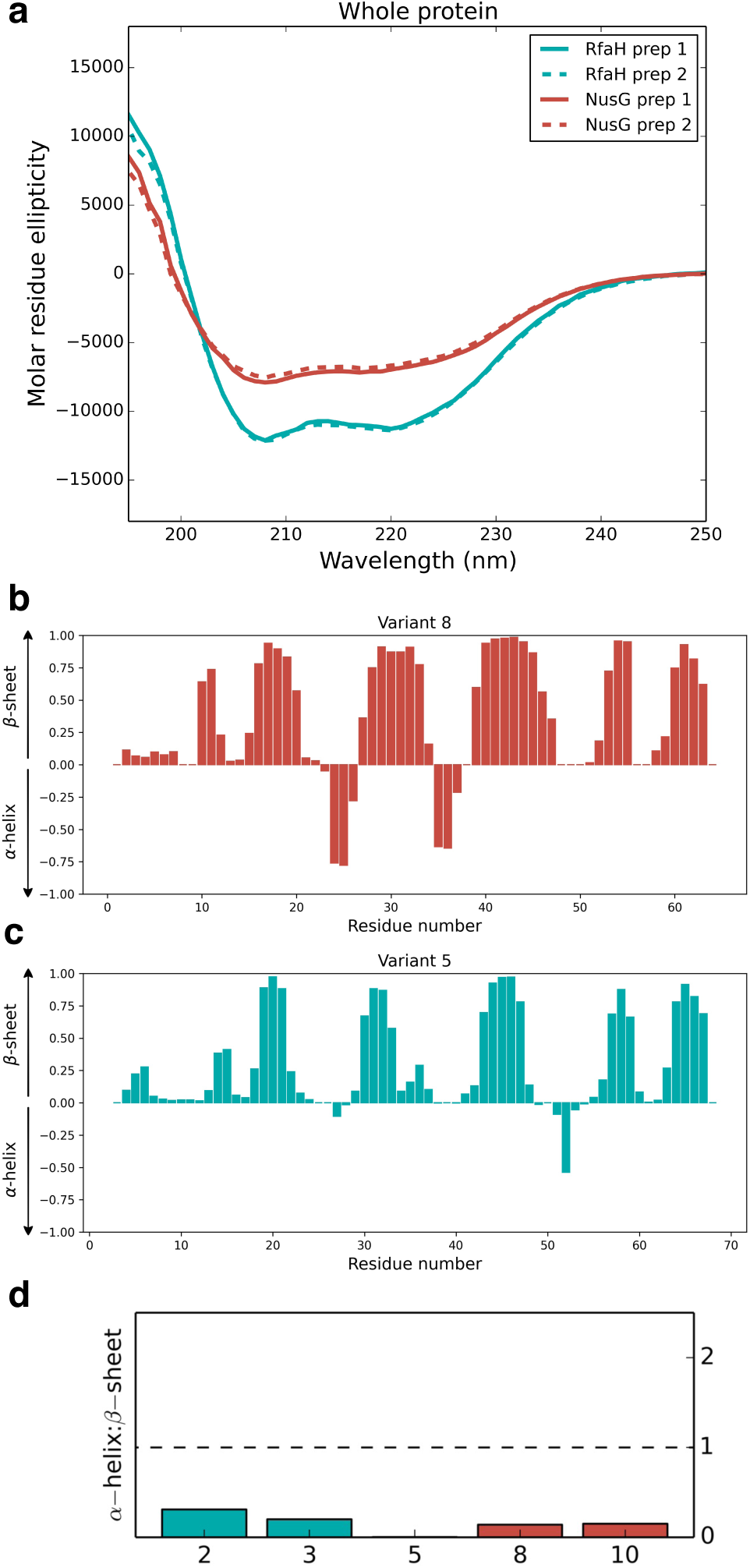
**(a)** Circular dichroism (CD) spectra of two different *E. coli* RfaH preps differ significantly from two different *E. coli* NusG preps. By contrast, CD spectra of the two different preps of both RfaH and NusG are nearly identical to one another. (**b-c**) TALOS+ secondary structure predictions of the assigned CTDs of Variant 8 (**b**) and Variant 5 (**c**). Plots suggest that both CTDs fold into structures with 5 β-sheets, consistent with the NusG β-roll fold. (**d**) CD spectra of 5 CTDs fold predominantly into β-sheets. All variants were estimated to have 27.3% (2)-36.2% (3) β-strand content, while α-helical content ranged from 0.03% (5)-8.4% (2). Like variants 2 and 3, variant 5 appears to switch folds. Secondary structure content was estimated by the BestSel server ^46^. Variant numbers correspond to those in **Fig. 2**.

**Extended Data Fig. 5.**
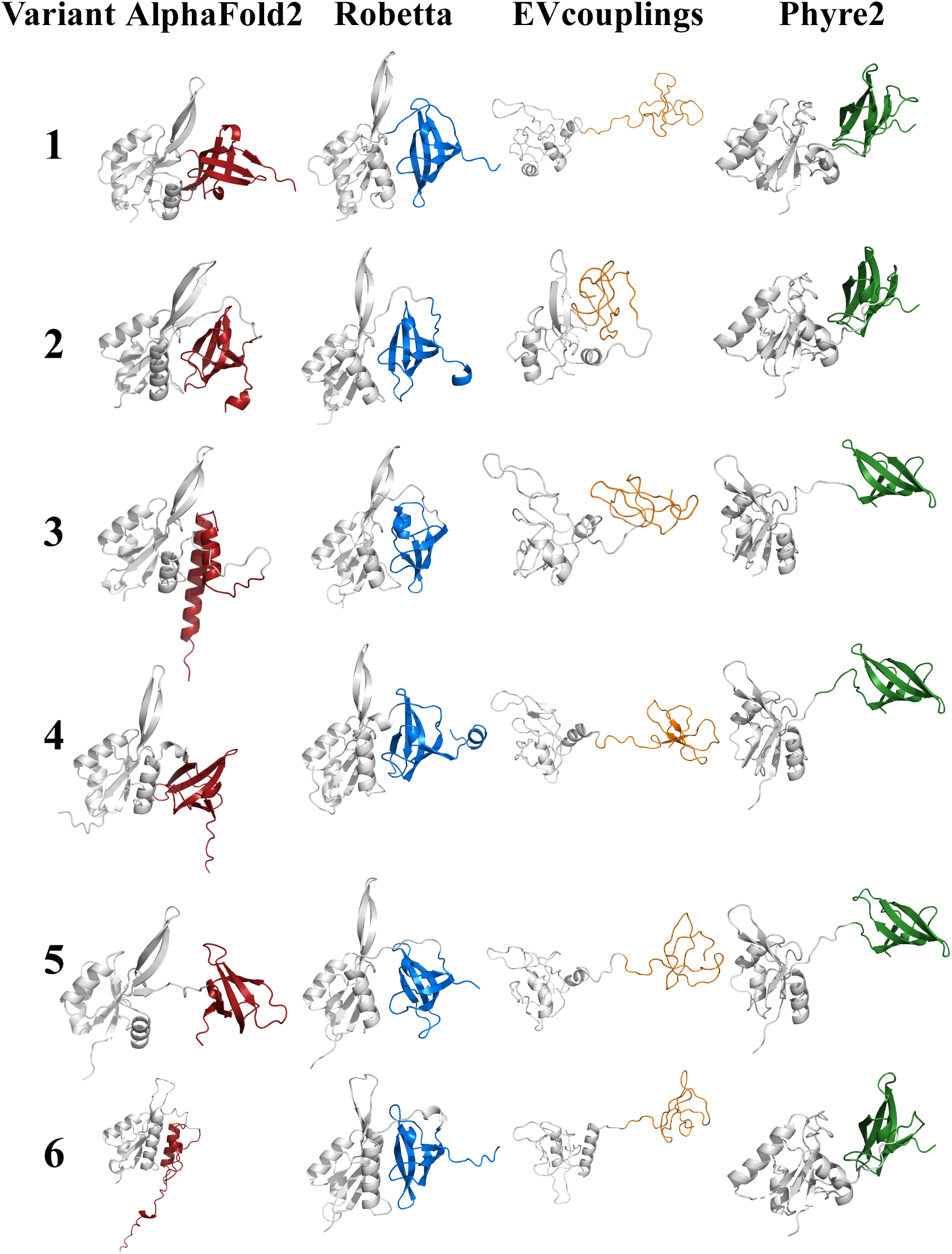
CTDs of the lowest-energy models for 6 proteins with RfaH-like folds (helical hairpin) are predicted assume β-sheet folds, including *E. coli* RfaH (Construct 3), which has an experimentally validated structure. CTDs are colored burgundy (AlphaFold2), blue (Robetta), orange (EVcouplings), green (Phyre2)

**Extended Data Fig. 6.**
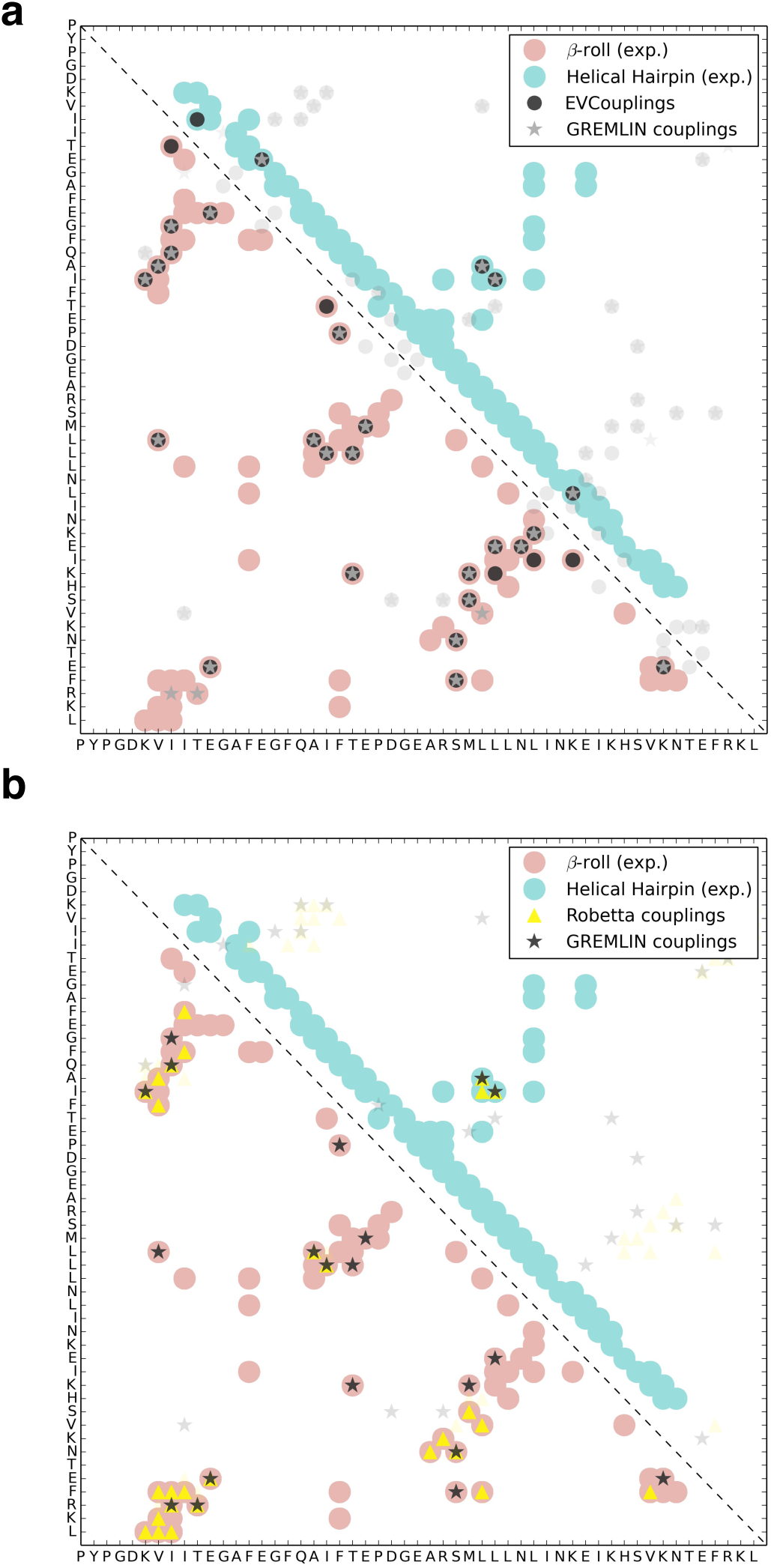
GREMLIN couplings calculated from EVCouplings (a) and Robetta (b) sequence alignments largely match contacts from the experimentally determined β-roll fold (red, PDB ID 2LCL) but did not match any contacts unique to the helical hairpin fold (teal, PDB ID 2OUG_A).

**Extended Data Fig. 7.**
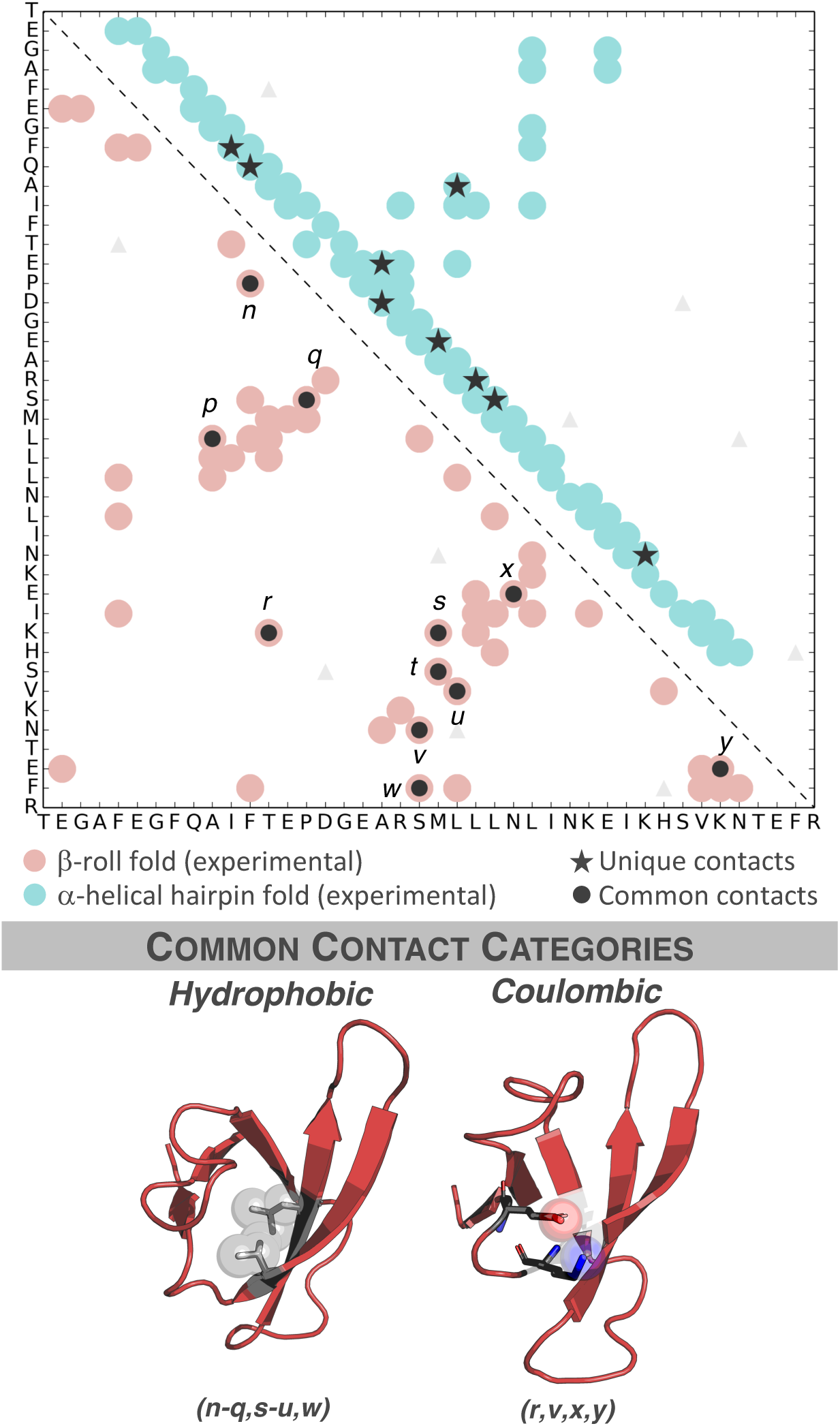
The single-fold paradigm biases protein structure predictions. EVCouplings and Robetta identify conserved residue-residue contacts (gray circles) corresponding to the β-roll fold of *E. coli* <u>RfaH</u> (PDB ID: 2LCL, red circles) but not the α-helical hairpin fold (PDB ID: 2OUG, teal circles). By contrast, secondary-structure-based predictions identified contacts from both the β-roll in common with EVCouplings and/or Robetta (gray circles) and the α-helical hairpin folds (gray stars). Faded triangular couplings do not correspond to either experimentally determined fold. Italicized letters correspond to individual contacts observed in the β-roll fold. Contact categories and their corresponding letters are shown below.

**Extended Data Fig. 8.**
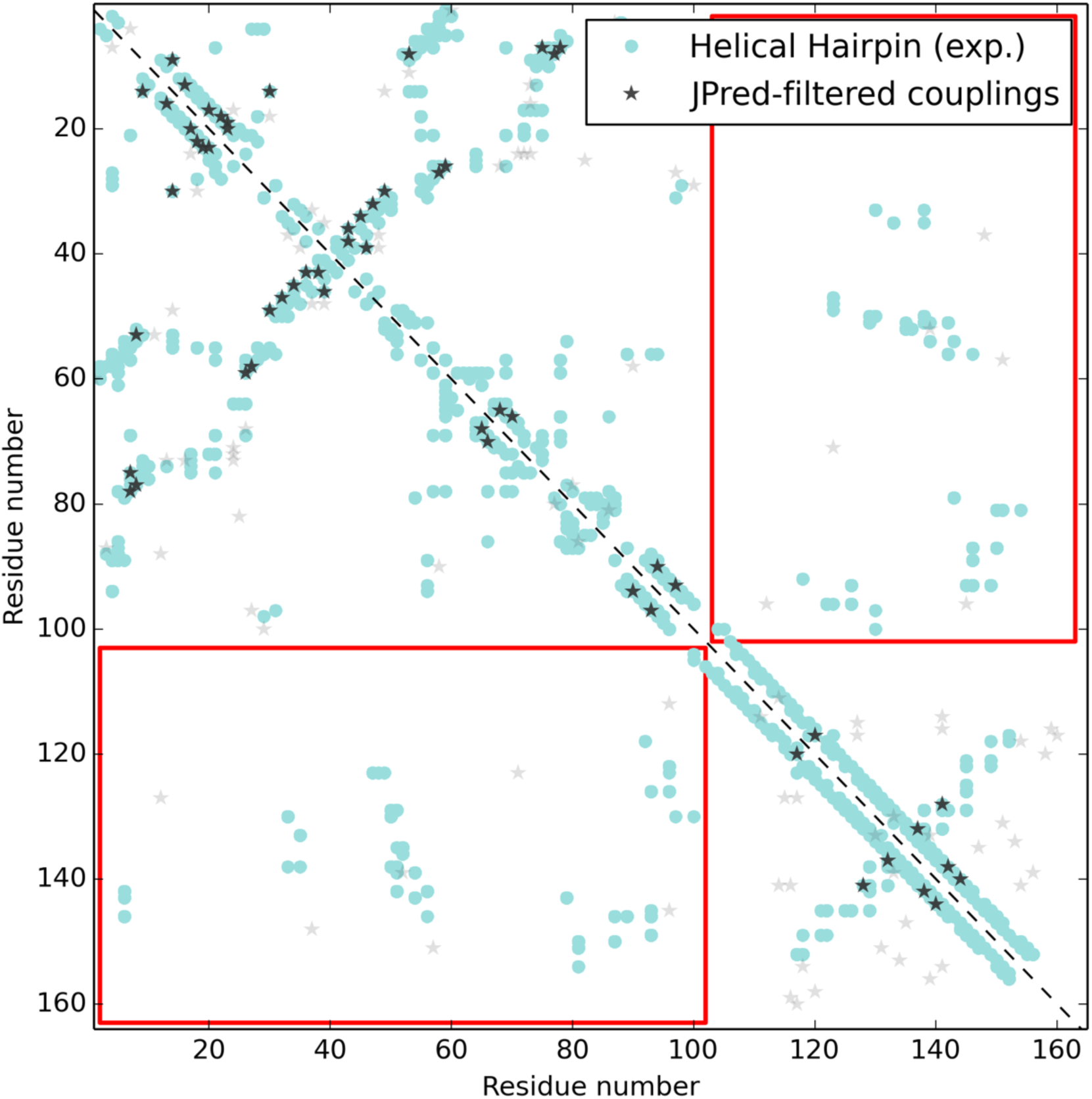
JPred-filtered couplings of putative full-length fold switchers calculated by GREMLIN. No interdomain contacts (found within red boxes) were consistent with the experimentally determined structure of full-length RfaH (PDB ID: 2OUG).

**Supplementary Table 1**. Annotations and predictions of all sequences identified in the NusG superfamily (Attached Separately).

**Supplementary Table 2.**
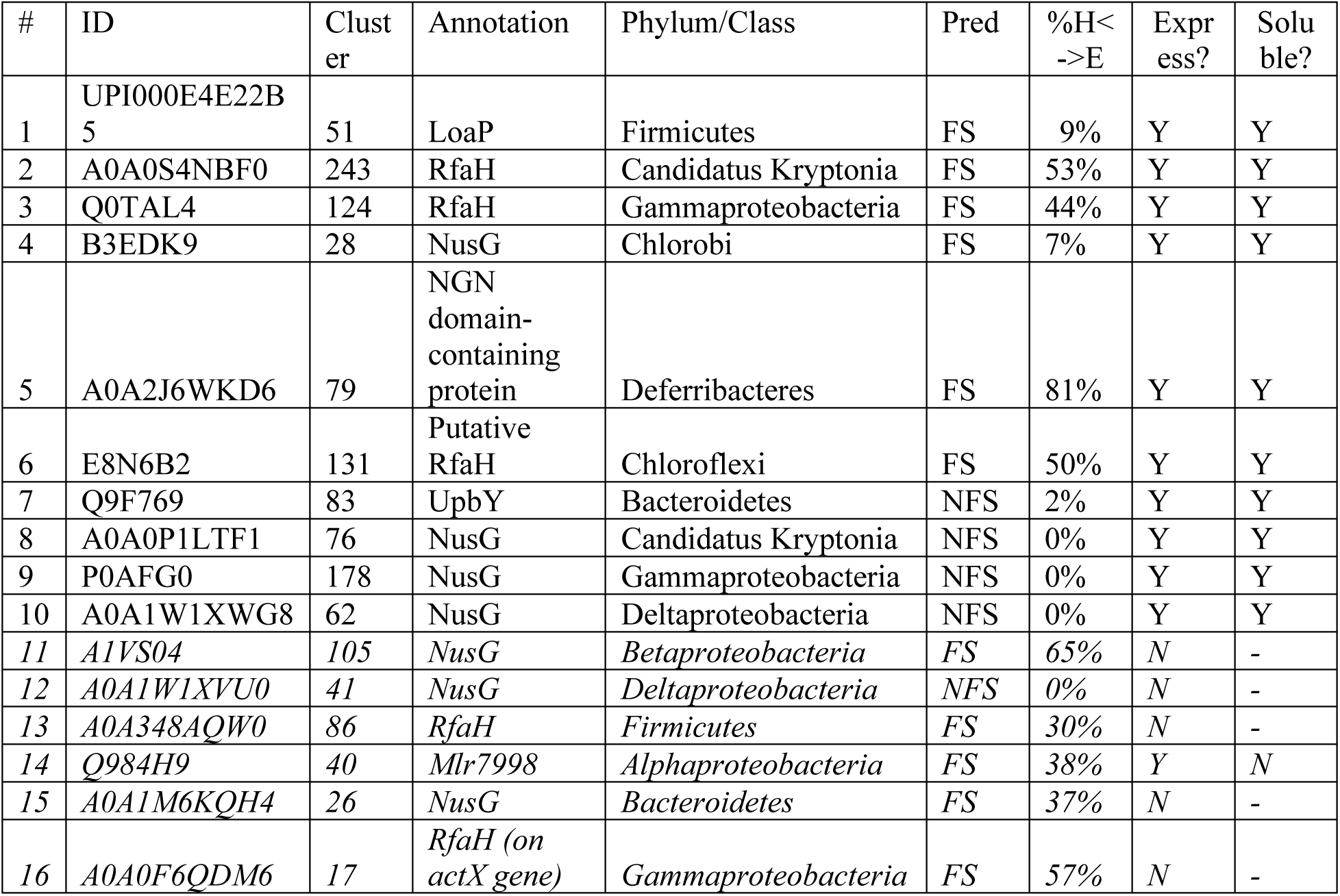
Sequences of all variants tested (see also Extended Data Figure 3).

**Supplementary Table 3.** Sequences of CTDs whose chemical shifts were assigned. See also Extended Data Figure 4.

| CTD Variant | Sequence |
| --- | --- |
| 8 | AELERIDVPF RVGDSVKVID GPFTDFSGVV QEVNSEKMKL<br>KVMINIFGRK TPVELDFTQV EIEK |
| 5 | MVDGFIDTKS EEFKKGDTIL IKDGPFKDFV GIFQEELDSK<br>GRVSILLKTL ALQPRITVDK DMIEKLHN |

**Supplementary Table 4**. Additional annotations and predictions of archaeal and eukaryotic sequences used to determine the tree in **Fig. 4**.

**Supplementary Table 5.**
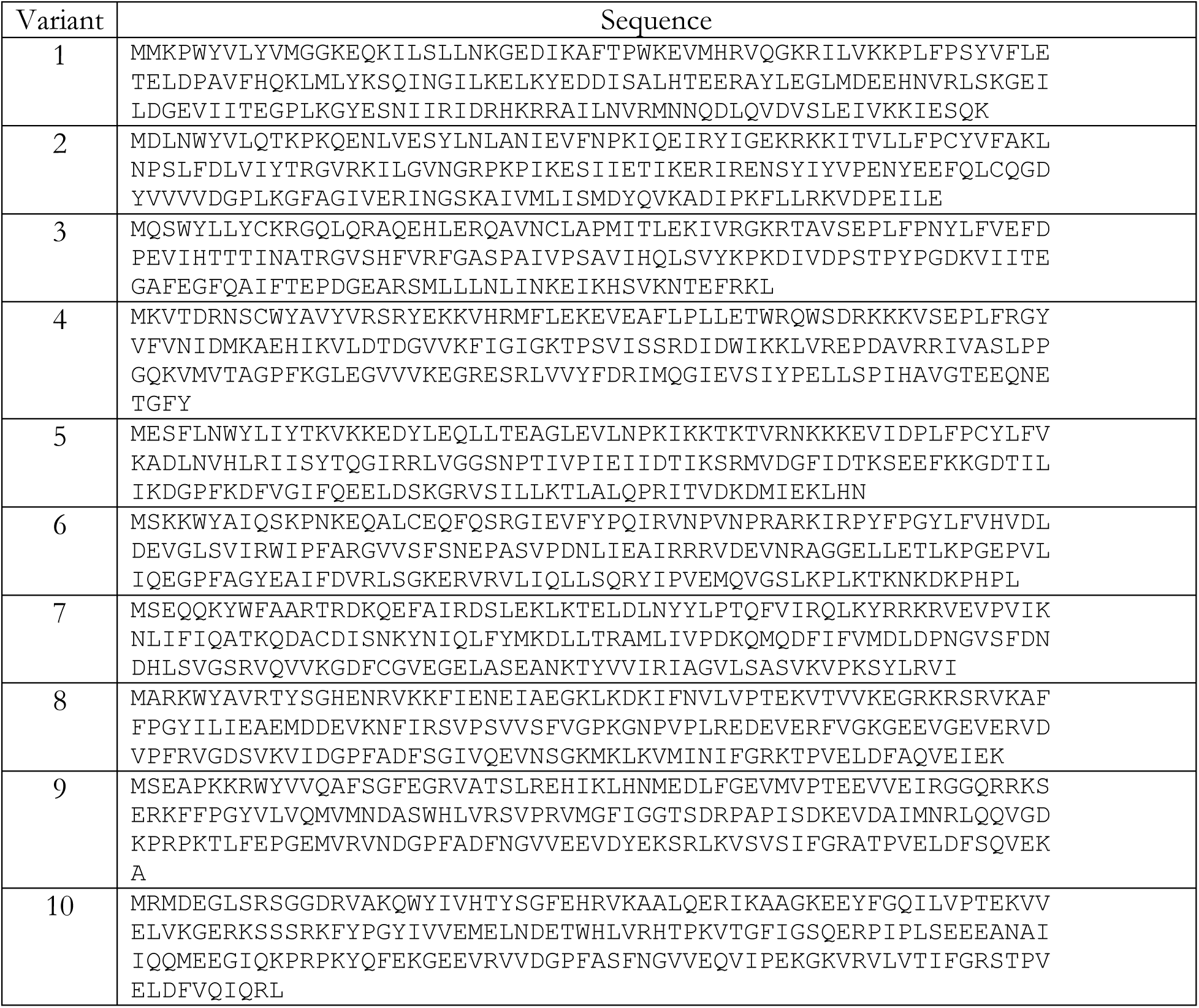
Sequences of all variants successfully purified and characterized by circular dichroism.

